# The StrainDiscoveryDatabase: an open framework for standardized microbial strain data

**DOI:** 10.64898/2026.09.09.750348

**Authors:** Julius F. Witte, Artur Lissin, Isabel Schober, Christian Ebeling, Helko Lüken, Julia Koblitz, Andrey Yurkov, Boyke Bunk, José Miguel López-Coronado, Adrian Vaello, Aurora Zuzuarregui, Rosa Aznar, Agustin García-Doñate, Ana M. P. Melo, Gerard Verkleij, Ronald P. de Vries, Vincent Robert, Wieland Meyer, Jörg Overmann, Lorenz C. Reimer

**Affiliations:** Leibniz Institute DSMZ – German Collection of Microorganisms and Cell Cultures; Spanish Type Culture Collection-University of Valencia (CECT-UVEG); Microbial Resource Research Infrastructure-Spanish node (MIRRI-ES); MIRRI-ERIC; Westerdijk Fungal Biodiversity Institute (WI-KNAW); SNSB- Staatliche Naturwissenschaftliche Sammlungen Bayerns

## Abstract

The vast amount of existing data on microbial strains holds immense potential to revolutionize bioindustry through the application of Artificial Intelligence (AI). However, the training of robust predictive AI models requires large-scale, unified, and non-redundant microbial datasets, which is currently severely hindered by the deep fragmentation of the data and the existence of synonymous strain identifiers in different culture collections. To overcome these infrastructural bottlenecks, we have established the *StrainDiscoveryDatabase* (SDD), a comprehensive, machine-readable dataset encompassing over 6.2 million harmonized data points for 256,889 microbial strains. The SDD does not rely on its own data repository, but rather on existing data that is retrieved on the fly from highly curated databases. Through an automated pipeline phenotypic, genotypic, and contextual data are systematically retrieved via the Application Programming Interfaces (APIs) of the Bacterial *Diversity* database (Bac*Dive*), the Microbial Resource Research Infrastructure Information System (MIRRI-IS) and the catalogue of the DSMZ. In order to reliably resolve synonymous strain identifiers, the StrainInfo database and its identification tools are employed, enabling the accurate deduplication and unification of records from disparate sources. The resulting aggregated, globally unique dataset is provided in a highly standardized JSON format in strict adherence to the FAIR data principles. By bridging isolated database silos and linking distributed knowledge to discrete biological entities, the SDD provides a high-quality, foundational resource designed to accelerate trait-based strain discovery, large-scale comparative analysis, and machine learning applications, thereby supporting the translation of the extensive existing knowledge on microbial traits to bioindustrial applications.

## Background & Summary

Microbial strains are the fundamental units of microbiological research, serving as the physical basis for novel discoveries in biotechnology, medicine, agriculture, ecology and beyond. Over the past decades, the increasing numbers of microbial strains and the growth in phenotypic and genomic data has resulted in an unprecedented accumulation of strain-specific information. To date the usability of this wealth of information is severely impeded by data fragmentation, redundancy and synonymous strain designations.

Information on growth conditions, metabolic capabilities, pathogenicity, and taxonomy is frequently siloed across culture collections and specialized databases. This hinders the trait-based discovery of microbial strains for bioindustrial and other applications. In particular, it limits the ability to perform large-scale, cross-dataset analyses such as the training of machine learning models or conducting comprehensive comparative genomics. So far, such analyses face significant logistical hurdles in data retrieval, standardization, and integration.

Several prominent data infrastructures have made substantial advances in aggregating and standardizing microbial strain data to overcome these bottlenecks. The Bacterial *Diversity* Database^1^ (Bac*Dive*), recently recognized as a Global Core Biodata Resource, provides highly curated phenotypic and genotypic information, with its recent 2025 iteration now encompassing 102,187 standardized strain records. At the same time, the Microbial Resource Research Infrastructure Information System^2^ (MIRRI-IS), connects researchers to high-quality biological resources and harmonized contextual data across European culture collections and currently provides data on 180,879 microbial strains. Despite their similar content and complementary nature, integrating data between such independently developed resources up to now remained challenging due to distinct data structures, ontologies, and access protocols. Furthermore, a single microbial strain is frequently deposited in multiple culture collections under varying designations (e.g., *Escherichia coli* K-12 has distinct accession numbers in the CECT, DSMZ, and BCCM collections). To aggregate the distributed knowledge for a particular strain, such as information from multiple culture collections, scientific publications and public database entries, a reliable method is needed to resolve the synonymous identifiers.

In this respect, the StrainInfo database^3^ serves as a critical centralized index that can be employed to overcome the complex challenge of synonymous identifiers. Recently re-developed as a core infrastructure by the NFDI4Microbiota consortium, StrainInfo systematically tracks microbial strains and their deposits across culture collections worldwide. By mapping equivalent strain designations and issuing persistent strain-level identifiers, StrainInfo provides the essential framework required to accurately link distributed data back to a single biological entity.

In this Data Descriptor, we present the *StrainDiscoveryDatabase* (SDD), a comprehensive, highly standardized, and machine-readable dataset encompassing integrated data for 256,889 microbial strains. We have developed a robust, automated pipeline that leverages the Application Programming Interfaces (APIs) of the Bac*Dive* and MIRRI databases, as well as the DSMZ catalogue to systematically retrieve raw strain data. We subsequently utilized the StrainInfo API to resolve synonymous strain identifiers, enabling the accurate deduplication and unification of records originating from disparate sources. The aggregated data was harmonized and structured into a standardized JSON format, ensuring strict adherence to the FAIR (Findable, Accessible, Interoperable, and Reusable) data principles^4^. The specific JSON format, termed the *Microbial Strain Data Standard* (https://github.com/LeibnizDSMZ/microbial-data-standard), is a product of the Horizon Europe project BioIndustry 4.0 (https://www.bioindustry4.eu). It was developed to store complex data from different sources in one file and thus enable the easy access and exchange of microbial strain data. By bridging the gap between isolated databases and resolving the complex web of strain synonymy, the SDD provides a unified, high-quality resource that will significantly accelerate data-driven discoveries and machine learning applications in biotechnology and microbiology in general.

The urgent demand for machine-readable microbial data is evidenced by recent specialized compilations, such as the BactoTraits dataset^5^, which incorporated 31 functional traits from Bac*Dive* for ecological community profiling. While such datasets provide high value for targeted applications like 16S rRNA biomonitoring, they rely on Bac*Dive* as the single data source, but do not address the broader infrastructural challenge of cross-database integration. To achieve a true and global harmonization between relevant datasets, it is necessary to go beyond single-database extraction. By integrating the MIRRI-IS API and leveraging StrainInfo to resolve complex identifier synonymy across multiple culture collections, the novel SDD provides a globally unique, FAIR-compliant resource at a significantly larger scale serving as a foundational architecture for predictive and comparative analysis.

## Methods

### Data acquisition and compilation

Three publicly available APIs were used as raw data sources for this dataset (Figure 1). The Bac*Dive* API (https://api.bacdive.dsmz.de/), the public version of the MIRRI-IS API (https://webservices.bio-aware.com/mirri_new/public) and the API of the DSMZ catalogue (https://api.strains.dsmz.de/docs). All APIs provide strain data under the Creative Common By Attribution license (CC-BY). Data retrieved from Bac*Dive* and MIRRI-IS were transformed into the *Microbial Strain Data Standard* using Python scripts. Data of the DSMZ catalogue API is already provided in the *Microbial Strain Data Standard* format.

From Bac*Dive* we requested 102,187 strains, while the MIRRI-IS API provided 180,879 accessible strains. Of these we were able to transform 99,537 strains from Bac*Dive* and 176,426 strains from MIRRI-IS into the standard. The remaining strains were discarded since the data did not fulfil the requirements defined by the *Microbial Strain Data Standard*, e.g. by missing essential taxonomic information (see the technical validation section for more detailed information). The DSMZ catalogue API provided data on 32,080 strains. After a deduplication step for each resource, 295,051 strains proceeded to the matching step (Bac*Dive*: 99,212 strains; MIRRI-IS: 163,928 strains; DSMZ catalogue: 31,911 strains).

**Figure #1:**
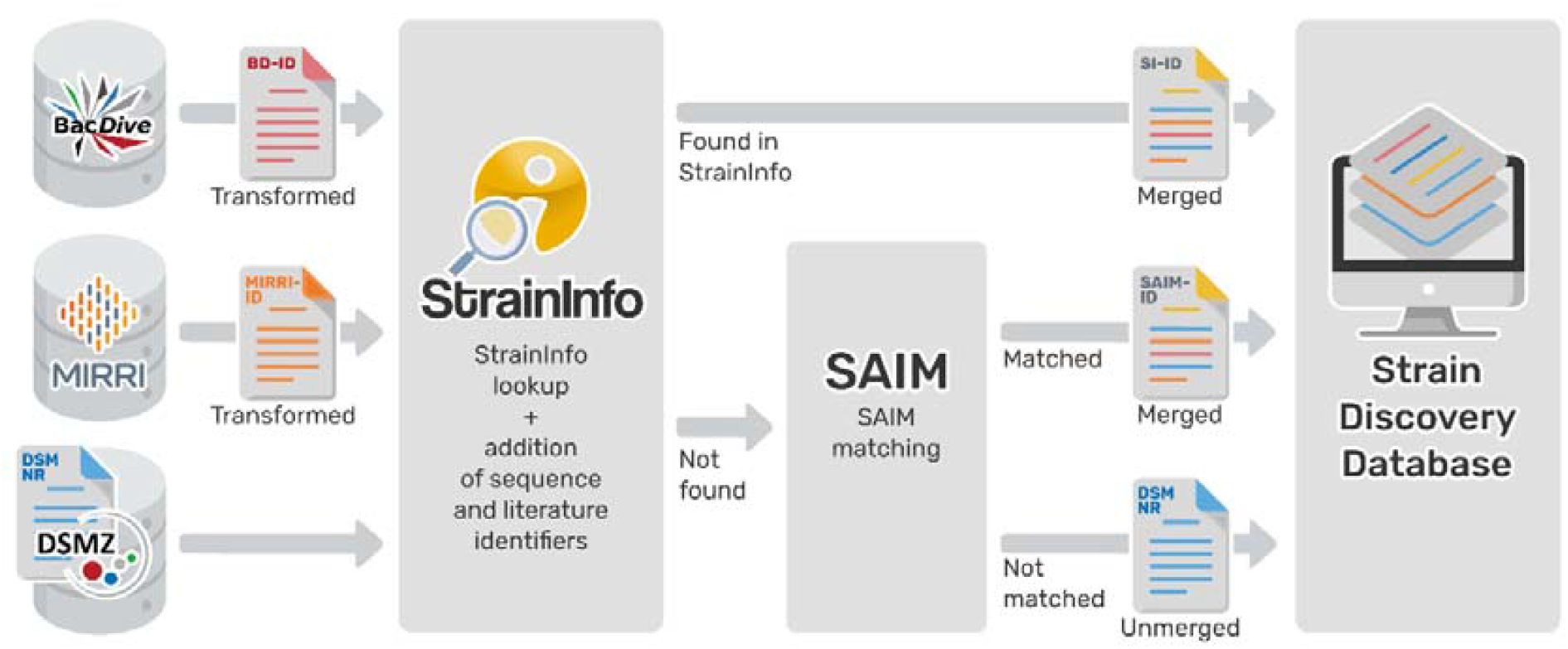
Overview of the data processing pipeline: Data is retrieved through the public APIs of the Bac*Dive* database, MIRRI-IS and the DSMZ catalogue. While DSMZ data is provided directly in the *Microbial Strain Data Standard* format, data from Bac*Dive* and MIRRI-IS has to be transformed. Through the StrainInfo API, strain synonymy is resolved and strains are merged accordingly. Strains unknown to StrainInfo are matched using the SAIM tool. The standardized and resolved strain data is integrated into the *StrainDiscoveryDatabase*.

### Matching of strains

The strain matching consists of two main consecutive steps. The first utilizes the StrainInfo-API (https://straininfo.dsmz.de/service) for strain identification, the second step is a resolution with the ‘Strain Authentication and Identification Methods’ (SAIM, https://github.com/LeibnizDSMZ/saim/^6^) library for all the strains which could not be identified by StrainInfo. The StrainInfo matching step first queries StrainInfo with the input strain culture collection numbers (CCNos) using the *v2/search/strain/cc_no* API endpoint, which returns StrainInfo IDs (SI-IDs) of all StrainInfo strains with hits. Next, details on all the gathered SI-IDs, including taxonomic information, are fetched via the *v2/data/strain/avg* endpoint, and each SI-ID is linked to an input CCNo through its corresponding StrainInfo deposit (represented by an SI-DP identifier). All the linked SI-IDs are then filtered based on their taxonomic compatibility with the input strain, first on the domain level, then on species or genus level. Finally, for each input strain, the SI-ID referenced by the majority of CCNos is chosen.

Strains that could not be matched in this way, are analyzed using the SAIM pipeline. SAIM looks for the largest overlaps in the CCNos; the procedure was described previously in Lissin *et al*.^3^. All SAIM matches are also filtered by their taxonomic information in the same way as described above. Every strain remaining unmatched retains its original primaryId as retrieved from the source database (e.g. Bac*Dive*, MIRRI-IS or DSMZ).

### Merging and checking the data

When a strain is identified by StrainInfo, additional data about this strain is retrieved from the StrainInfo database and added to the dataset. The integrated information encompasses strain origin, sequence accession numbers, scientific literature, collection information and strain identifiers, as well as taxonomy and type strain information.

Whenever two input strains are identified as describing the same strain, they are merged, retaining all input data. All data points are referenced by a source which provides a direct link to the respective data entry in the original database. All data that is stored in lists as objects is combined, while the source and relatedData links are updated. RelatedData represents a relationship between multiple data fields of data objects, e.g. a growthCondition relation between testTemperature and cultivationMedia. Whenever two data objects are identical except for their sources, the data will appear only once in the dataset but is referenced by both sources to avoid data duplication. Strain data files that are not identified by StrainInfo, are processed using the SAIM tool and are merged if possible.

## Data Record

### Dataset description

The data set described here comprises harmonized data on 256,889 strains and enables to find strains beyond the barriers of single databases and catalogues. The scripts provided utilize the respective APIs to integrate data from Bac*Dive* (99,212 strains), MIRRI-IS (163,928 strains), and the DSMZ catalogue (31,911 strains) (Figure 2). Bac*Dive* and MIRRI-IS each themselves provide aggregated data from many culture collections. The merged strain data is made available through the SDD in a standardized and machine interpretable way.

**Figure #2:**
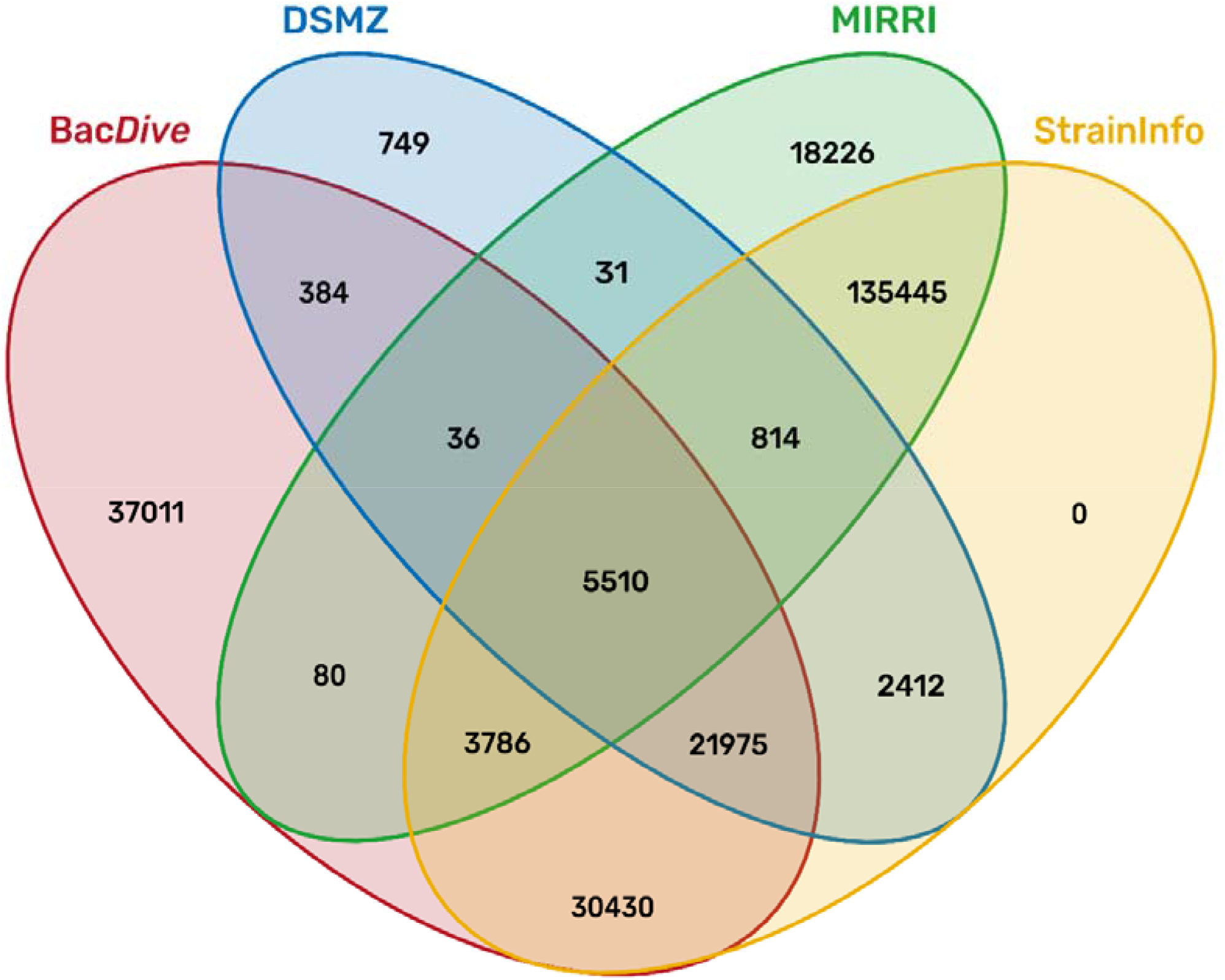
Overview of the distribution of 256,889 matched strains: Distribution across all four data sources after strain matching. StrainInfo adds no additional strains, as it was used only to resolve the matching of strains provided by the other databases. 200,372 strains could be resolved using the StrainInfo database (yellow ellipse). 934 additional strains not found in StrainInfo could be matched using the SAIM tool and have been merged to 531 strains in the final dataset (all areas that overlap but are not inside the yellow ellipse).

So far, the data set comprises 24 standardized data fields for information relevant for the selection of strains for research and industrial applications (Table 1). They include taxonomy, cultivation, metabolic functions, origin, safety and legal information as well as important links to sequence and literature information. The number of strains for which a data field is filled ranges from only 1-2 % e.g. for known applications and pathogenicity up to 80-100% e.g. for origin, growth conditions and taxonomy. Documentation and examples for all data fields are provided in Table 1.

**Table #1:** Data field overview: *Total* = total number of entries in the dataset; *Strains filled* = number of strains where this data field is filled; *Coverage %* = percentage of strains where this data field is filled;

| Field | Total | Strains filled | Coverage % | Description |
| --- | --- | --- | --- | --- |
| bioSafety | 232,513 | 198,724 | 77.4 | Biosafety classification of the strain. e.g. BSL 1/2 |
| collections | 368,096 | 221,764 | 86.3 | In which culture collections the strain is available. |
| cultivationMedia | 332,324 | 183,532 | 71.4 | Cultivation media the strain grows on. e.g. marine agar (MA) |
| enzymes | 382,176 | 29,106 | 11.3 | Enzyme activity of the strain. e.g. urease positive |
| fattyAcidProfiles | 5,831 | 5,571 | 2.2 | The fatty acid profile of the strain. The fatty acids present in the cells and their shares in the complete fatty acid content. |
| gcContent | 24,662 | 15,985 | 6.2 | The GC content of the genome of a strain. |
| growthConditions | 272,488 | 209,682 | 81.6 | Conditions the strain needs to grow, including temperature, pH and oxygen relation. |
| halophily | 9,627 | 9,573 | 3.7 | The halophily or halotolerance of the strain. |
| hemolysis | 816 | 814 | 0.3 | The type of hemolysis the strain is capable of and the blood type it has been tested on (if available). |
| identifier | 1,249,093 | 256,889 | 100 | Culture collection numbers, designations, and database identifiers related to a strain. |
| knownApplications | 3,221 | 3,008 | 1.2 | Known application for the strain, e.g. photoproduction of hydrogen. |
| legal | 183,872 | 173,278 | 67.5 | Legal information and restrictions on the use of the strain, e.g. Nagoya restrictions or dual use. |
| literature | 446,347 | 32,299 | 12.6 | Scientific literature associated with the strain. |
| metabolites | 1,187,075 | 36,937 | 14.4 | Metabolites produced or utilized by the strain, e.g. glucose. |
| morphology | 24,453 | 22,157 | 8.6 | Cell and colony morphology of the strain, e.g. cell shape, flagellum arrangement, colony colour. |
| origin | 476,780 | 226,794 | 88.3 | Origin of the strain, including sampling and isolation data. |
| pathogenicity | 11,584 | 5,145 | 2 | Whether the strain is pathogenic including human, plants and animals. |
| relatedData | 32,080 | 31,911 | 12.4 | Link that defines the relationship between two or more data fields whose conditions depend on each other, e.g. temperature and cultivation medium. |
| sequences | 171,339 | 49,513 | 19.3 | Sequence accessions for nucleotide or protein sequences of a strain. |
| sporeFormation | 5,706 | 5,687 | 2.2 | Whether the strain is able to produce spores and what kind. |
| staining | 21,465 | 21,357 | 8.3 | Results of staining tests performed for the strain, including Gram staining. |
| taxon | 510,038 | 255,445 | 99.4 | Taxonomy of the strain. |
| tolerances | 38,410 | 8,505 | 3.3 | The tolerance levels to chemical substances, e.g. antibiotics, the strain exhibits. |
| typeStrain | 241,028 | 239,131 | 93.1 | Whether a strain is the type strain of the species it belongs to. |

In total 6,231,024 data points are currently collected in the SDD, covering microbial strain data across the domains of Bacteria (52.57%), Archaea (0.47%), Fungi (46.28%), Algae (0.68%) and Protists (0.002%) (Table 2).

**Table #2:**
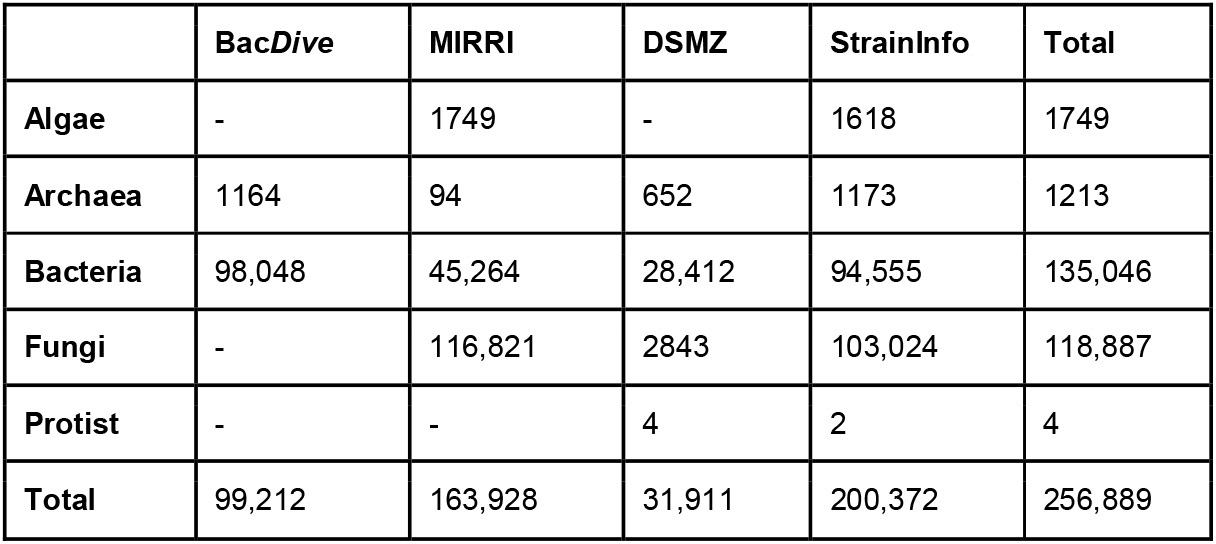
Organism types per source after matching.

|  | BacDive | MIRRI | DSMZ | StrainInfo | Total |
| --- | --- | --- | --- | --- | --- |
| Algae | - | 1749 | - | 1618 | 1749 |
| Archaea | 1164 | 94 | 652 | 1173 | 1213 |
| Bacteria | 98,048 | 45,264 | 28,412 | 94,555 | 135,046 |
| Fungi | - | 116,821 | 2843 | 103,024 | 118,887 |
| Protist | - | - | 4 | 2 | 4 |
| Total | 99,212 | 163,928 | 31,911 | 200,372 | 256,889 |

## Technical Validation

The *Microbial Strain Data Standard* is defined and publicly available on GitHub (https://github.com/LeibnizDSMZ/microbial-data-standard). Data from the Bac*Dive* and MIRRI-IS APIs is transformed into the format and verified using the provided Pydantic model, which functions as a validator and guarantees technical correctness.

The most common validation issue arising during this step is missing or unknown organism type information, as this field is required by the model and any strain with an unclear organism type will be rejected. For Bac*Dive* this error type occurs for 503 strains and for MIRRI-IS for 3619 strains. In 48 strains, issues with malformed literature URLs, origin tags (MISO), geographic locations, malformed date formats (Sampling, Isolation) and further occurred. The corresponding strains were discarded.

## Usage Notes (Outlook and future usage)

The *StrainDiscoveryDatabase* marks a milestone in the handling of microbial strain data by providing the first open framework that enables researchers to generate up-to-date, standardized and FAIR datasets on microbial strain knowledge, that can be fully integrated into their own workflows and easily expanded using their own tools and data.

By utilizing public web services of large strain databases to integrate up-to-date information, combined with the power of the *Microbial Strain Data Standard* to ensure comparability, this framework opens up new possibilities for leveraging strain data in both research and industry. To give a few examples: the database can be queried to find microbial strains growing under specific conditions, having distinct metabolic capabilities or underlying certain legal requirements. Notably, the dataset can also be used as machine interpretable input for complex analyses like using AI to make genome-based predictions where experimental data is missing^7^. It is important to note that the dataset presented here is only a starting point: the source databases are constantly being expanded and further data resources providing additional information on existing and new strains can easily be integrated, so that this database can quickly grow. Users can contribute their own code and own data, which will further increase the huge potential for future usage.

Finally, this dataset provides a deep insight into the heterogeneity of data from different strain resource collections (Figure 3). It underlines the importance of a central data format in view of gathering, matching and comparing strains. It also shows the feasibility and usability of the *Microbial Strain Data Standard* as well as the possibilities its widespread adoption could offer for the exchange, comparison, and analysis of microbial strain data. The provision of special new API endpoints offering data of databases and culture collections using the *Microbial Strain Data Standard*, for example, could save users the time and effort needed to make non-standardized data from different sources interoperable.

**Figure #3:**
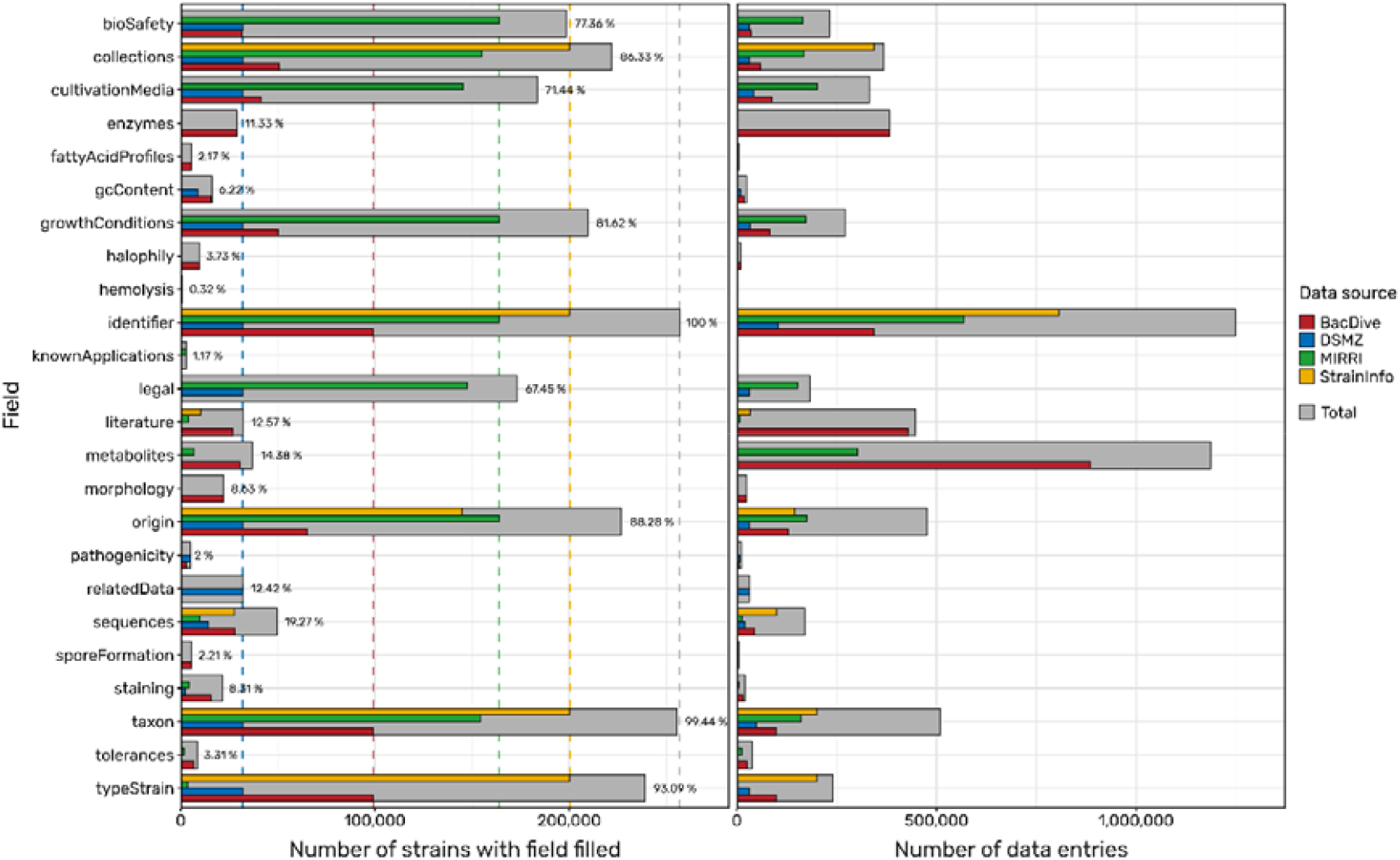
Data field statistics: Bar plots showing for how many strains a data field is filled (left) and the number of data entries per data field (right). Grey bars represent total numbers for the data set, coloured bars represent entries contributed by the respective source database. Dashed lines in the left panel mark 100% of total strains in the data set (grey line) and 100% of strains contributed by each data source (coloured lines). Due to the performed strain merging and dereplication of field entries, the number of total strains/entries are less than the sum of the source strains/entries.

## Data Availability

All data described in this study is publicly available and published under the Creative Commons BY Attribution license (CC-BY). Data is available through Zenodo (https://doi.org/10.5281/zenodo.21379409) or can be recreated via the pipeline hosted on GitHub (https://leibnizdsmz.github.io/strain-discovery-database/)

## Code Availability

All code described in this study is publicly available and published under the MIT license. Repository holding the pipeline to retrieve, transform and match the data: https://leibnizdsmz.github.io/strain-discovery-database/ (https://doi.org/10.5281/zenodo.21375665)

Other repositories related to the project: https://github.com/LeibnizDSMZ/microbial-data-standard (https://doi.org/10.5281/zenodo.21374532) https://github.com/LeibnizDSMZ/saim (https://doi.org/10.5281/zenodo.14879790)

